# TheCellVision.org repository: expansion with high-content cell imaging projects on eukaryotic intracellular organization and DUB biology

**DOI:** 10.64898/2026.09.02.748561

**Authors:** Myra Paz David Masinas, Athanasios Litsios, Matej Usaj, Harsha Garadi Suresh, Charles Boone, Brenda J. Andrews

## Abstract

High-content cell imaging approaches enable the systematic characterization of cellular function through the acquisition of multimodal information from large cohorts of live single cells. Yet, due to their scale and complexity, data acquired via such approaches are often challenging to meaningfully share across laboratories and effectively use for independent studies. Since its inception, the main purpose of TheCellVision.org repository has been to fill this gap, providing the research community with access to large-scale, multimodal single-cell datasets, in a structured, intuitive, and user-friendly way. Here, we report on the third major update of TheCellVision.org, which involves the expansion of the repository with the addition of data from two single-cell phenomics projects; the Intracellular Organization Dynamics project, which quantitatively maps changes in the morphology of 21 major subcellular structures in live yeast cells elicited by the systematic inhibition of essential genes, and the DUB Biology project, which describes changes in the concentration and localization of the budding yeast proteome in mutants of key deubiquitination enzymes (DUBs). With these additions, the repository now hosts six complementary high-content imaging projects which collectively explore the dynamics of intracellular organization and the proteome during changes in cell state and in response to environmental and genetic perturbations.

**ARTICLE SUMMARY:** TheCellVision.org is a repository for visualizing and mining data from yeast high-content imaging projects in an intuitive way. We report the 3^rd^ major expansion of TheCellVision.org, which involves the addition of two new projects; the Intracellular organization dynamics project, which quantitatively describes morphological changes in intracellular organization upon systematic inhibition of essential genes and progression of cells towards death, and the DUB Biology project, which describes genome-scale changes in protein localization and concentration in key deubiquitylase mutants of budding yeast.

## INTRODUCTION

High-content screening (HCS) methodologies enable the image-based phenotypic profiling of millions of live cells in different genetic backgrounds and across diverse environmental conditions. In *Saccharomyces cerevisiae (budding yeast)*, this is usually achieved using systematic genetics for the construction of large arrays of strains expressing desired fluorescent reporters and/or carrying mutations of interest, the subsequent use of high-throughput (HTP) live-cell fluorescence microscopy for imaging cells of the respective strains, and finally, the application of computational tools for the extraction and analysis of multiparametric phenotypic features from single cells.

Due to their comprehensive nature and multi-dimensionality, HCS datasets can often be repurposed to address aspects of cell biology different from (or complementary to) those for which they were originally generated. As such, images derived from HCS projects can serve as rich resources for future analysis by the scientific community (see for example (Huh et al. 2003)). Yet, due to their scale and complexity, meaningfully disseminating HSC datasets among laboratories and other end-users for independent analysis poses a major challenge. For example, our recent HCS project generated 129,525 microscopy images and other complementary genome-scale datasets of different modalities, and involved the extraction and analysis of multi-parametric information from >20 million live cells and >4,000 unique proteins (Litsios et al. 2024).

TheCellVision.org is an online repository for hosting image data from HCS yeast projects in a structured way, enabling users to intuitively access, visualize, and download the respective datasets. In the last database update (Masinas et al. 2024), we described four HCS datasets housed in TheCellVision.org (CYCLoPS; Endocytic Compartment Morphology; Cell Cycle Omics; PIFiA) exploring diverse aspects of yeast cell biology. Here, we report on the 3^rd^ major update of TheCellVision.org, which involves its expansion to include image datasets from two new HCS projects: [1] the Intracellular Organization Dynamics project, which explores morphological changes elicited in 21 organelles and major subcellular structures upon inhibition of essential genes and progression of cells towards death (Litsios et al. 2026), and [2] the DUB Biology project, which describes changes in the concentration and localization of the yeast proteome caused by mutation of key deubiquitination enzymes (DUBs) (Garadi Suresh et al. 2024). With these additions, the repository now hosts images from six complementary high-content imaging projects (**Figure 1**) which collectively explore the dynamics of intracellular organization and the proteome during diverse developmental cellular programs as well in response to various environmental and genetic perturbations.

**Figure 1.**
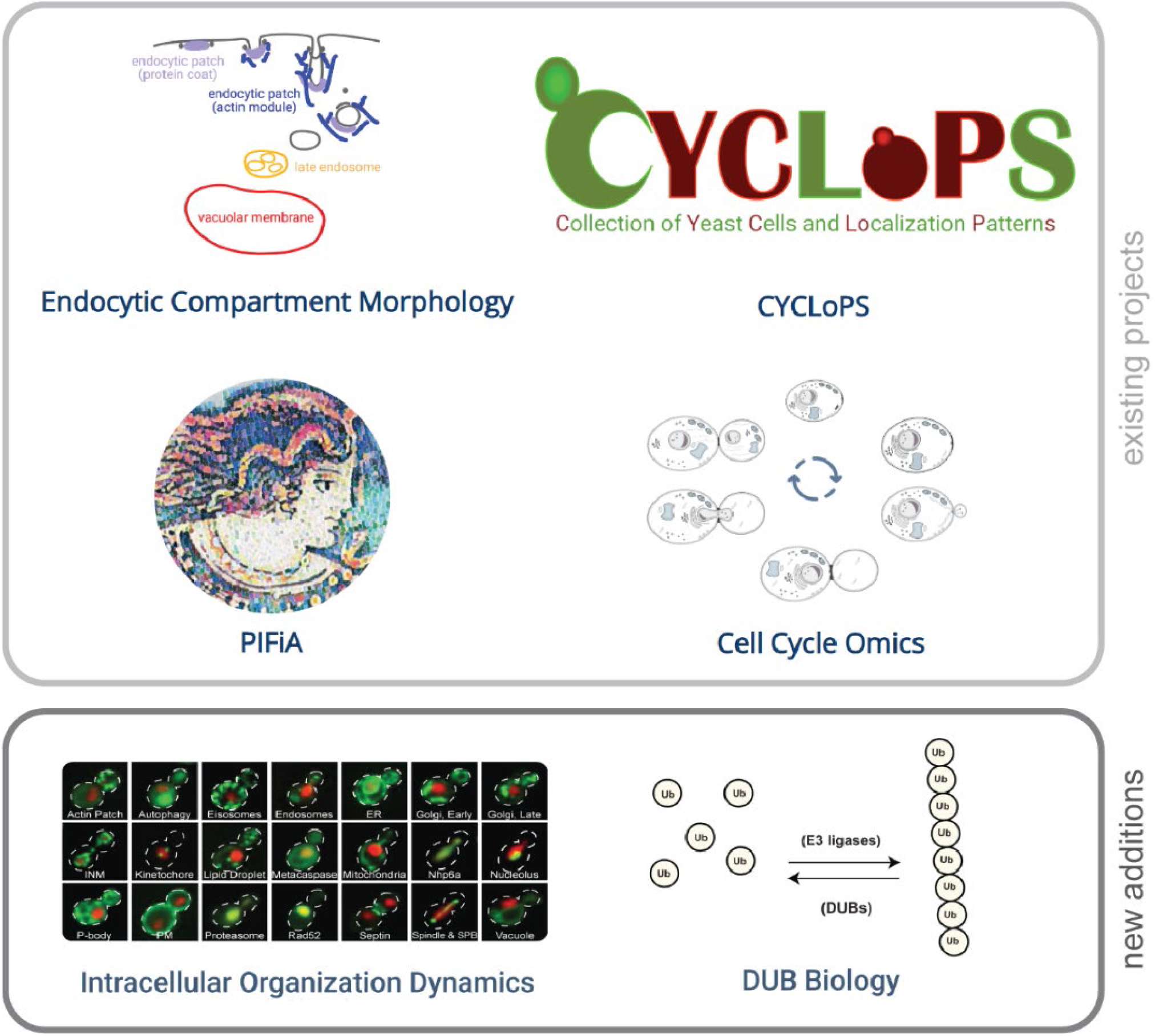
Overview of HCS projects housed in TheCellVision.org. Snapshots from the landing page of TheCellVision.org showing currently hosted projects. Existing and newly added projects are annotated.

## MATERIALS AND METHODS

### High-throughput imaging screens

Prior to this update, TheCellVision.org housed data from four large-scale projects (Masinas et al. 2020;Masinas et al. 2024) and two new datasets have been added (**Figure 1**):

#### Previously available datasets

i. CYCLoPs (*Collection of Yeast Cell and Localization Patterns*) (Chong et al. 2015; Koh et al. 2015; Kraus et al. 2017).
ii. Endocytic Compartment Morphology (Mattiazzi Usaj et al. 2020)
iii. Cell Cycle Omics (Litsios et al. 2024).
iv. PIFiA (*Protein Image-based Functional Annotation*) (Razdaibiedina et al. 2024)

#### Newly added datasets

v Intracellular Organization Dynamics. This is a newly added project that describes changes in intracellular organization elicited via the systematic inhibition of essential genes and progression of cells towards death(Litsios et al. 2026). Individual organelles and other major subcellular structures were visualized via fusion of native yeast proteins with GFP. Temporal resolution was achieved via time-lapse live-ciell fluorescence confocal imaging. The function of essential genes was perturbed using temperature-sensitive (TS) alleles (Li et al. 2011).
vi DUB Biology. This project describes changes in the abundance and subcellular location of the yeast proteome elicited in single and double KO mutants of key DUBs (Garadi Suresh et al. 2024). Changes in protein abundance and localization were quantified via imaging of cells expressing native yeast proteins fused with GFP at their endogenous loci, and the application of a convolutional neural network for image analysis.

More information about (i-ii) and (iii-iv) can be found in (Masinas et al. 2020)and (Masinas et al. 2024), respectively.

### Databased development and schema

We use PostgreSQL, a relational database management system, to store and organize the data in TheCellVision.org. The database schema has been expanded to include two new main clusters – one for each of the new research projects: Intracellular Organization Dynamics and DUB Biology (**Figure S1**).

## RESULTS AND DISCUSSION

### INTRACELLULAR ORGANIZATION DYNAMICS

In the Intracellular Organization Dynamics project, we used high-content screening to explore how intracellular organization changes when the function of essential genes is perturbed (Litsios et al. 2026). The status of intracellular organization was assayed by monitoring the morphology of 21 organelles and other major subcellular structures through the endogenous fusion of one of their native proteins with C-terminal GFP tags (Huh et al. 2003). Essential gene function was perturbed using temperature-sensitive (ts) alleles (Li et al. 2011); this ts strain collection included 352 essential genes spanning all fundamental cellular processes. In total, ∼7,400 yeast strains – each representing a different combination of a subcellular structure monitored and an essential gene perturbed – were shifted to the restrictive temperature and were imaged every 3h for a total of 24h using live-cell HTP confocal microscopy.

This resulted in the collection of 1,732,344 microscopy images across all fields of view and channels which contained data from ∼70.5 million single cells. To identify which of these cells displayed morphologically abnormal subcellular structures, single-cell features were extracted and a machine learning framework for outlier detection based on One-Class Support Vector Machines (Schölkopf et al. 1999; Mattiazzi Usaj et al. 2020) was applied. The final analyzed dataset contained a time-resolved quantitative account of morphological defects elicited in intracellular organization upon inhibition of essential yeast genes, expressed via *penetrance*, i.e. the percentage of cells in the population that showed morphological defects in each of the monitored subcellular structures, for each of the perturbed essential genes, and for each of the timepoints assayed.

Upon selecting the Intracellular Organization Dynamics project at TheCellVision.org, the user is directed to the project’s main page (**Figure 2A**) where they can retrieve information about the related publication (‘Publication’), download its supplemental material (‘Supplemental files’), review the list of genes that were assayed (‘Gene List’), and access a description of the resource alongside information on how to use it (‘About’). On this page, the user is prompted to select a gene of interest from the dropdown menu or enter the name of the gene directly into the query box. When available, names of human orthologs are also indicated in the dropdown menu (in parentheses next to the yeast gene name), and they can also be directly typed into the query box. Upon selection, the user is then navigated to the page containing the results related to the perturbation of the respective gene (**Figures 2B-2D)**.

**Figure 2.**
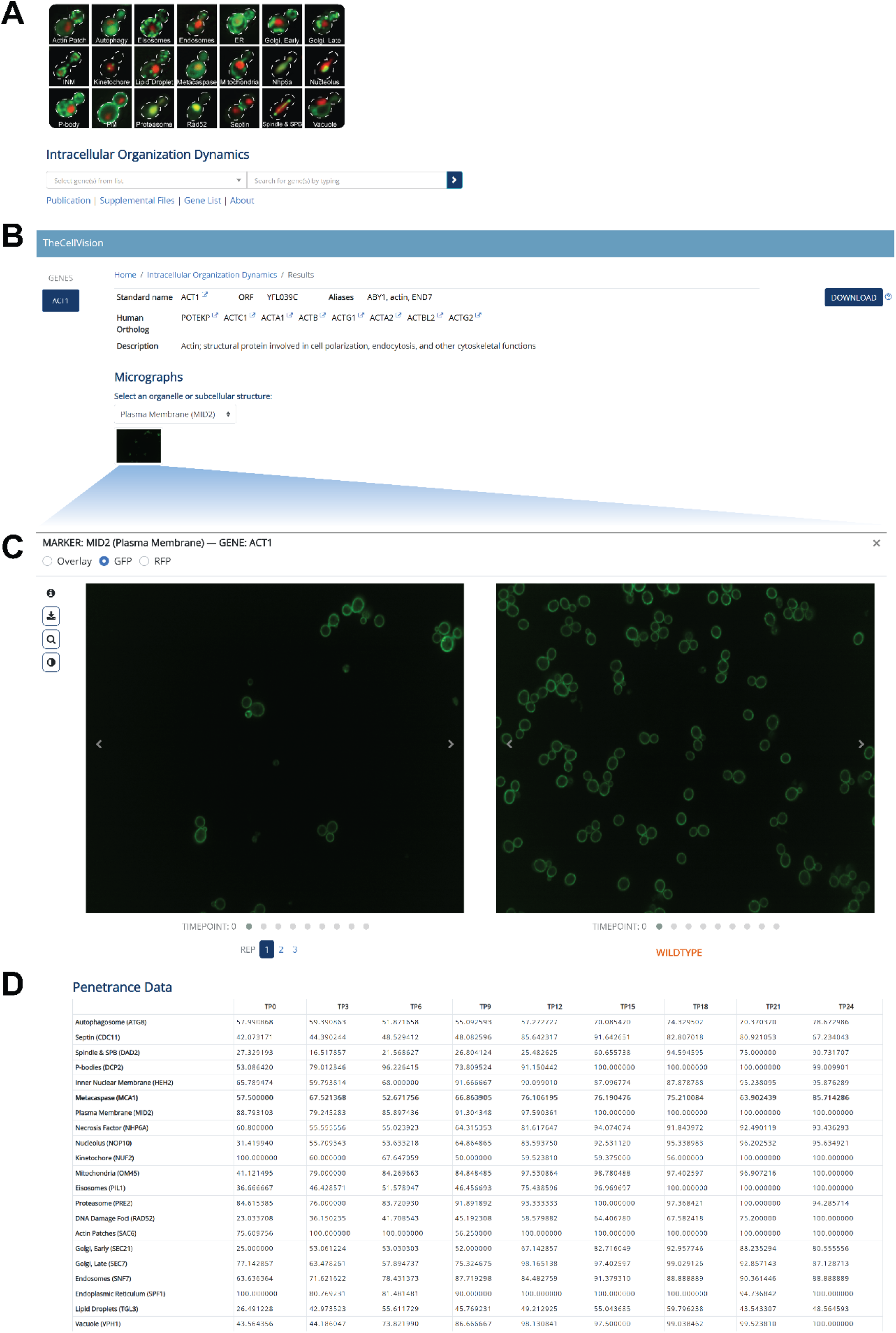
Layout overview of Intracellular Organization Dynamics main and results pages. (A) Snapshot of main page of Intracellular Organization Dynamics project, displaying the indicated functions and gene search prompts. (B-D) Snapshots of various parts of the results section after the search for an example gene that was perturbed (*ACT1* in this case). (B) Overview of gene name(s) and functional description. (C) Cell viewer for examination of microscopy images. In this specific example, images from the Mid2 (Plasma Membrane) marker are shown. (D) Penetrance values (percentage of cells with morphologically abnormal structure) at each time point for each of the subcellular structures monitored.

Here, information about the assayed gene is displayed, along with a brief description about its function (**Figure 2B**). At the upper right side of this page, the user can select to download results associated with the query gene in the Intracellular Organization Dynamics project (‘DOWNLOAD’ function). Underneath the gene description is the *Micrographs* module, where microscopy images of single cells are displayed across all timepoints and available biological replicates (**Figure 2C**). Here, the user is prompted to select an organelle or subcellular structure from the dropdown menu, and upon opening the cell viewer, they can browse microscopy images of single cells in the mutant (left) and wild-type control (right) backgrounds. The subcellular structure is displayed in green (GFP channel) and a nuclear marker (same across all strains) in red (RFP channel). The user can select to view specific channels individually or an overlay of them, as well as to adjust the brightness of images and magnify regions of interest within them. Furthermore, the vertical arrow at the left of the cell viewer allows users to directly download the microscopy images.

Finally, underneath the cell viewer, the analyzed data of the project are displayed in tabular format. Each row represents a different subcellular structure, and each column the penetrance of the assayed gene for this structure at the indicated timepoint (**Figure 2D**).

### DUB Biology

The DUB Biology project contains quantitative information related to changes in the abundance and subcellular localization of the yeast proteome in key single and double KO DUB mutants (Garadi Suresh et al. 2024). Changes in the proteome were assessed by analysis of single-cell images of the yeast protein-GFP collection – a resource of >4,000 strains each of which expresses a different protein tagged C-terminally with GFP at its endogenous locus. Three versions of the protein-GFP collection were constructed, each in a different genetic background: [1] *ubp2Δ*, [2] *ubp14Δ*, and [3] *ubp2Δ ubp14Δ*. All these collections, together with the *wild-type* control array, were imaged using HTP confocal microscopy, and single-cell microscopy images were analyzed using DeepLoc (Kraus et al. 2017; Litsios et al. 2024), a convolutional neural network for automated prediction of protein localization with resolution of 21 subcellular classes. Upon analysis of the 442,366 microscopy images and 9,626,924 single-cell crops, the final dataset consisted of protein abundance measurements based on mean GFP intensity (Mean ± Standard Deviation across all cells within a specific genetic background) for each of the proteins in each of the four collections. Moreover, for each protein in each of the four collections a vector representing the distribution of the protein over the 21 localization classes was produced.

Similarly to the Intracellular Organization Dynamics project, the landing page of the DUB Biology project (https://thecellvision.org/dubbiology/) allows the user to review the related publication, download its supplemental material, and retrieve information about the resource and how to use it (**Figure 3A**). Upon entering a gene/protein name of interest into the query box, the user is again navigated to the relevant results page that includes information about the gene and a data-download function (**Figure 2B**), underneath which the *Micrographs* module with the cell viewer can be found (**Figure 3C**). Here, the protein encoded by the queried gene is displayed in green (GFP channel), and a nuclear and cytoplasmic marker (same across all strains), in red (RFP channel) and blue (FarRed channel) respectively. Microscopy images from two biological replicates are displayed side by side, with the option to switch between the various screens performed (WT, and single and double KO mutants). The rest of the functionalities remain the same as is in the cell viewer of the Intracellular Organization Dynamics project.

**Figure 3.**
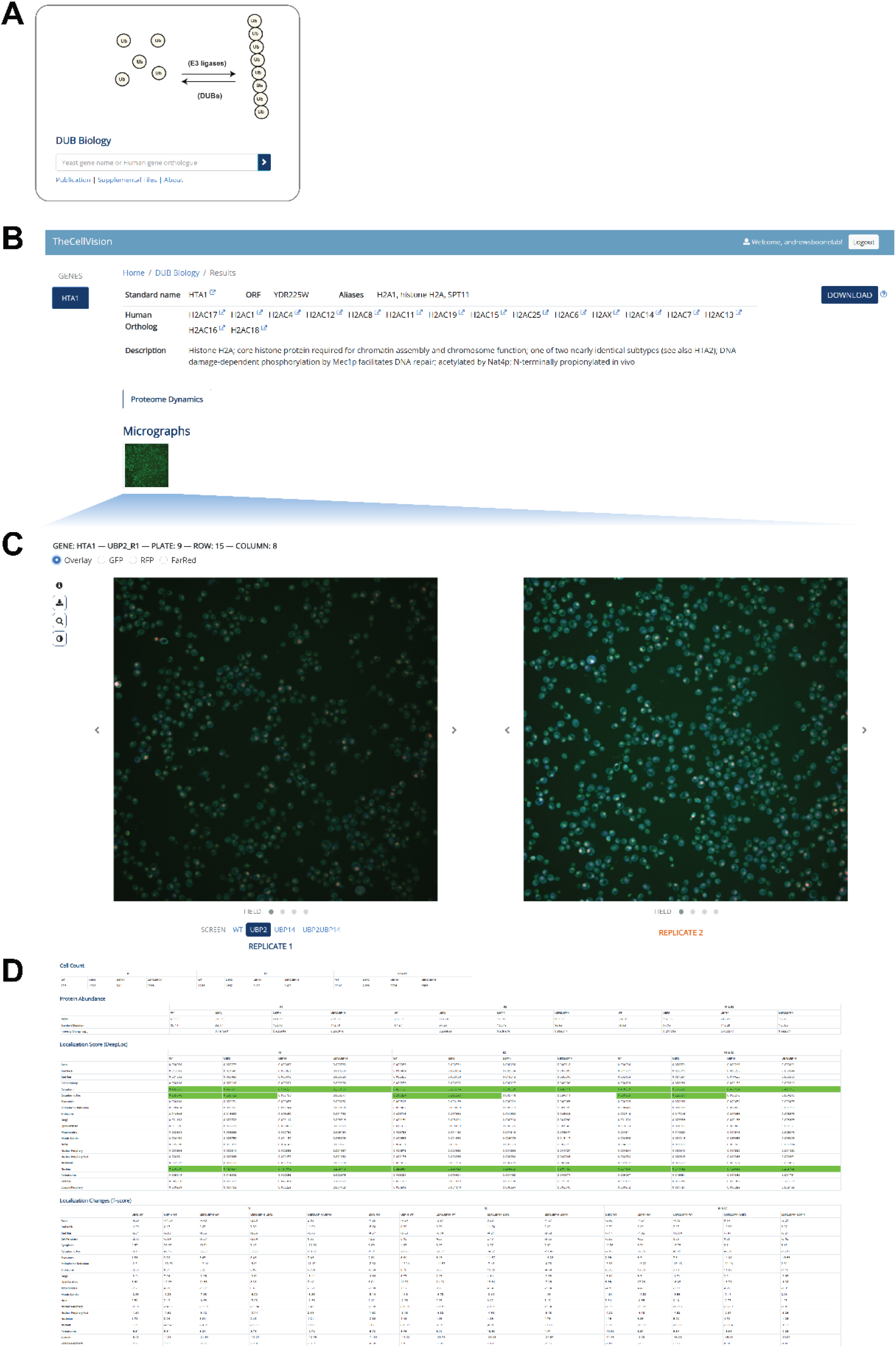
Layout overview of DUB Biology main and results pages. (A) Snapshot of main page of DUB Biology project, displaying the indicated functions and the gene search prompt. (B-D) Snapshots of various parts of the results section after the search for an example gene/protein (*HTA1* in this case), the dynamics of which were monitored in the various screens. (B) Overview of gene name(s) and functional description. (C) Cell viewer for examination of microscopy images. (D) Related results from the analysis of the single-cell images.

Underneath the cell viewer, quantified information from the microscopy images is presented (**Figure 3D**). This includes [1] the number of single cells analyzed in each replicate and screen for the queried gene/protein (*Cell Count*), [2] summary statistics regarding the abundance of the protein (*Protein Abundance*), [3] the subcellular localization of the protein with resolution of 21 localization classes (*Localization Score*; values highlighted in green denote significant localization scores), and [4] quantification of changes in the localization of the protein among any two genetic backgrounds screened (*Localization Changes*; values highlighted in blue and yellow denote significantly negative and positive localization changes, respectively). In all cases, information is presented for each individual biological replicate, as well as for data from both replicates combined.

## CONCLUSIONS AND FUTURE DIRECTIONS

Decoding cellular function at a systems-level entails understanding how molecular changes and their interactions translate to higher-order cellular phenotypes. High-content screening (HCS) methodologies provide a platform for dissecting such complex relationships, by enabling the multiparametric, image-based phenotypic profiling of millions of live cells in different genetic backgrounds and in diverse environmental conditions. Nevertheless, the challenge posed by the scale and complexity of such projects in accessibly storing and disseminating the resultant datasets can substantially limit their impact on scientific discovery.

TheCellVision.org serves as a user-friendly, central repository for live-cell fluorescence microscopy images and associated quantitative data generated using HCS approaches in budding yeast. By hosting raw and analyzed data in a structured and intuitive way, and providing the tools necessary for their visualization, exploration, and download, it facilitates the re-usability of these resources from independent researchers. The utility of TheCellVision.org as a platform for the exploration of new hypotheses, the independent re-analyses of existing large-scale dataset, and dataset benchmarking, is reflected in its continually increasing adoption by the scientific community; as of August 2026, the repository has had 49490 unique users from 120 countries.

TheCellVision.org will continue to expand and is scheduled to soon be supplemented with an additional large-scale phenotypic screen which provides a comprehensive morphological analysis of 18 subcellular yeast compartments upon genome-wide perturbations.

## DATA AVAILABILITY

Microscopy images, datasets, and computational tools associated with the previous projects are available as described in (Masinas et al. 2020; Masinas et al. 2024). Raw images associated with DUB Biology are available for download at https://www.ebi.ac.uk/biostudies/bioimages/studies/S-BIAD1164. Bulk download of images from Intracellular Organization Dynamics can be provided upon request. Source code and usage examples for DUB Biology are available at https://github.com/BooneAndrewsLab/DUB_Biology. The code for Intracellular Organization Dynamics pipeline is available at https://github.com/BooneAndrewsLab/Intracellular_Organization_Dynamics.

## SUPPLEMENTAL MATERIAL

**Figure S1.**
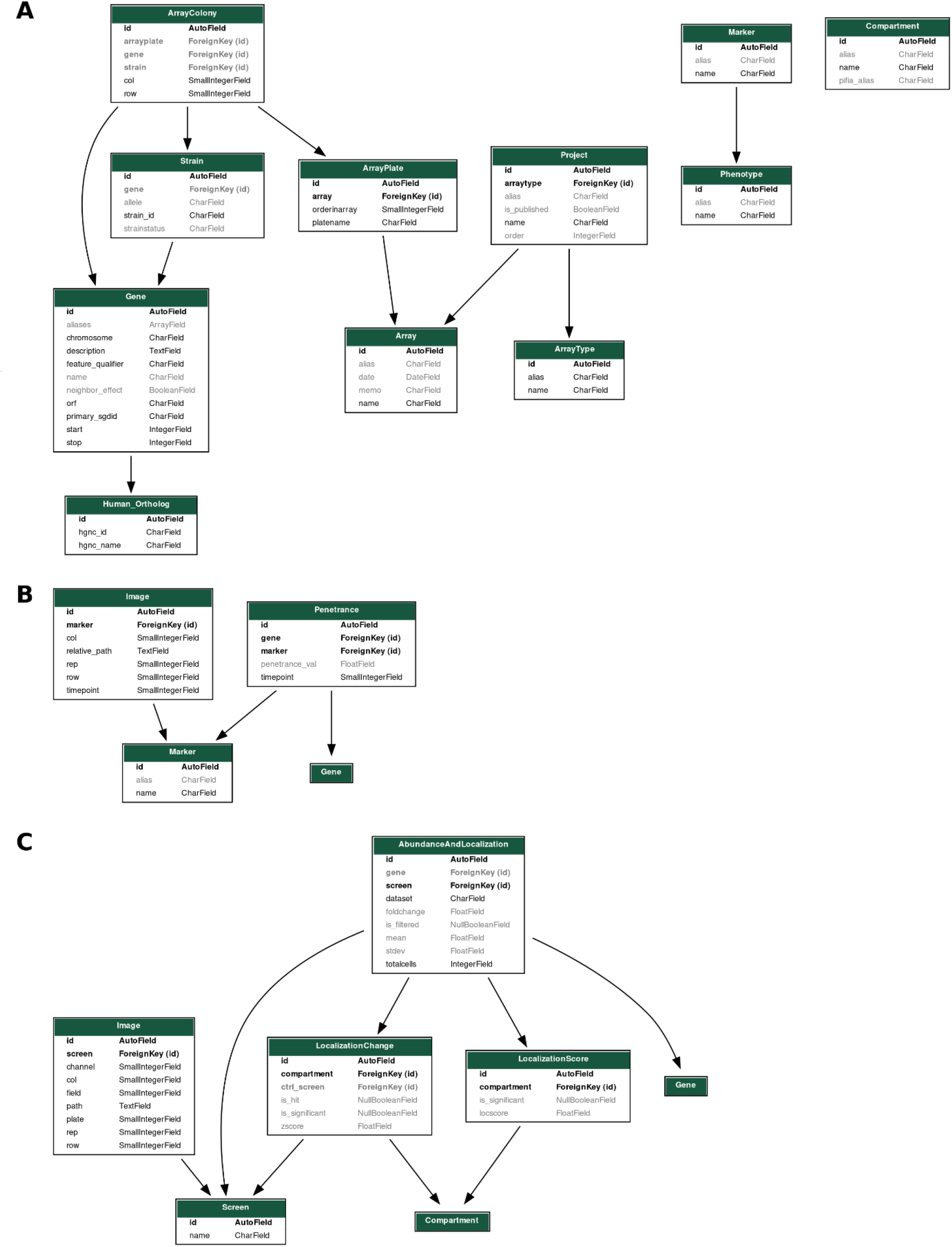
Database schema. Diagrams illustrating the database schema for the updated core cluster and two new projects: (A) core, (B) Intracellular Organization Dynamics project, and (C) DUB Biology project.

## REFERENCES

Chong YT et al. 2015. Yeast proteome dynamics from single cell imaging and automated analysis. Cell. 161(6):1413–1424 [accessed 2019 Oct 17]. 10.1016/j.cell.2015.04.051

Garadi Suresh H et al. 2024. K29-linked free polyubiquitin chains affect ribosome biogenesis and direct ribosomal proteins to the intranuclear quality control compartment. Mol Cell. 84(12):2337–2352.e9. 10.1016/j.molcel.2024.05.018

Huh W-K et al. 2003. Global analysis of protein localization in budding yeast. Nature. 425(6959):686–91. 10.1038/nature02026

Koh JLY et al. 2015. CYCLoPs: A Comprehensive Database Constructed from Automated Analysis of Protein Abundance and Subcellular Localization Patterns in Saccharomyces cerevisiae. G3 (Bethesda). 5(6):1223–32. 10.1534/g3.115.017830

Kraus OZ et al. 2017. Automated analysis of high-content microscopy data with deep learning. Mol Syst Biol. 13(4):924 [accessed 2020 Oct 19]. https://pubmed.ncbi.nlm.nih.gov/28420678/. 10.15252/msb.20177551

Li Z et al. 2011. Systematic exploration of essential yeast gene function with temperature-sensitive mutants. Nat Biotechnol. 29(4):361–367 [accessed 2022 Nov 9]. https://pubmed.ncbi.nlm.nih.gov/21441928/. 10.1038/NBT.1832

Litsios A et al. 2024. Proteome-scale movements and compartment connectivity during the eukaryotic cell cycle. Cell. 187(6):1490–1507.e21. 10.1016/j.cell.2024.02.014

Litsios A et al. 2026. Organelle interdependencies underlie the collapse of eukaryotic intracellular organization during cell death and aging. bioRxiv. 2026.08.25.746993 http://biorxiv.org/content/early/2026/08/28/2026.08.25.746993.abstract. 10.64898/2026.08.25.746993

Masinas MPD et al. 2020. TheCellVision.org: A Database for Visualizing and Mining High-Content Cell Imaging Projects. G3 (Bethesda). 10(11):3969–3976 [accessed 2023 Jun 19]. https://pubmed.ncbi.nlm.nih.gov/32934016/. 10.1534/G3.120.401570

Masinas MPD et al. 2024. Expanding TheCellVision.org: a central repository for visualizing and mining high-content cell imaging projects. Genetics. 227(1). 10.1093/genetics/iyae044

Mattiazzi Usaj M et al. 2020. Systematic genetics and single-cell imaging reveal widespread morphological pleiotropy and cell-to-cell variability. Mol Syst Biol. 16(2) [accessed 2022 Nov 4]. https://pubmed.ncbi.nlm.nih.gov/32064787/. 10.15252/MSB.20199243

Razdaibiedina A et al. 2024. PIFiA: self-supervised approach for protein functional annotation from single-cell imaging data. Mol Syst Biol. 20(5):4. 10.1038/s44320-024-00029-6

Schölkopf B et al. 1999. Support Vector Method for Novelty Detection. In: Solla S, Leen T, Müller K, editors. Advances in Neural Information Processing Systems. Vol. 12. MIT Press https://proceedings.neurips.cc/paper_files/paper/1999/file/8725fb777f25776ffa9076e44fcfd776-Paper.pdf

